# The *Candida* Genome Database

**DOI:** 10.64898/2026.07.28.741317

**Authors:** Jodi Lew-Smith, Shuai Weng, Gavin Sherlock

## Abstract

The *Candida* Genome Database (CGD; www.candidagenome.org) is both a model organism database and a fungal pathogen database. As a model organism database, CGD stores data for *Candida albicans*, which serves as a model species both for other *Candida* spp. and for non-*Candida* fungi that form biofilms and undergo routine morphogenic switching. As a fungal pathogen database, CGD now hosts locus pages for six species of the best-studied pathogenic fungi in the *Candida* group. Pathogenic *Candida* species have become increasingly drug resistant and there is thus a pressing need for research into basic *Candida* biology, epidemiology, phylogeny, and potential new antifungals, as well as a single location where all of the available data are collected, curated, and made easily searchable. CGD curates the gene-based *Candida* experimental literature in real time, extracting, organizing and standardizing gene annotations. CGD also links clinical data on disease to relevant Literature Topics to improve searchability for clinical researchers. Because CGD curates the literature for multiple species and most research focuses on aspects related to pathogenicity, we focus our curation efforts on assigning Literature Topic tags, collecting detailed mutant phenotype data, and assigning controlled Gene Ontology terms with accompanying evidence codes. Our Summary pages for each locus include the primary name and all aliases for that locus, a description of the gene and/or gene product, detailed ortholog information with links, a synteny view, a JBrowse window with a visual view of the gene on its chromosome, links to Phenotype, Gene Ontology, Interactions, and Expression pages, as well as sequence information, references cited on the summary page itself, and any locus notes. The database also serves as a community hub, where we link to various types of reference material of relevance to *Candida* researchers, including colleague information, news, and notice of upcoming meetings. We routinely survey the community to learn how the field is evolving and how needs may have changed. Here we describe CGD’s new modern web interface and multiple new tools that have been added in the last 6 months, allowing, among other things, users to better understand the available expression data for a locus and seamlessly switch between species for a given locus.

## Introduction

The Candida Genome Database currently annotates research data for six pathogenic *Candida* species, adding new ones when a sufficient community need exists. The current species are *C. albicans, C. glabrata* (*Nakaseomyces glabratus*), *C. dubliniensis, C. parapsilosis, C. auris* (*Candidozyma auris*), and *C. tropicalis*. Note, the genus *Candida* is a large, highly polyphyletic group of budding, white colony-forming yeasts in the subphylum Saccharomycotina, originally grouped together because of their similar morphology and lack of a defined teleomorph (sexual reproductive stage) and many *Candida* species have been recently renamed to better reflect their phylogeny (see Kidd et al. 2023), though this is not without controversy (see Denning 2024b). Following consultation with the community, CGD still uses the *Candida* species names. Among the species that CGD curates, *C. albicans* is by far the best studied of these pathogens and serves as a model for studying the less experimentally tractable members of the group (Kabir et al. 2012).

*C. albicans* infection in both immunocompetent and immunocompromised patients can result in several common diseases, including playing a role in dental caries as well as mucosal candidiasis of the oral cavity and the genitourinary tract (Eidt et al. 2020; Giri and Kindo 2012; Lass-Florl et al. 2024; Pfaller and Diekema 2007; Schelenz 2008; Vila et al. 2020). However, in immunocompromised patients, invasive candidiasis can lead to very high mortality rates, exceeding 35% (Bays et al. 2024; Denning 2024a; Lass-Florl et al. 2024). Furthermore, *Candida albicans* is the third or fourth most common nosocomial bloodstream isolate and not only is treatment costly but infection significantly extends the length of hospitalization (e.g., Andes et al. 2012; Dodds Ashley et al. 2012; Moran et al. 2009; Wan Ismail et al. 2020).

The majority of invasive Candida infections are caused by *C. albicans, C. glabrata, C. parapsilosis, C. tropicalis, C. krusei*, and, more recently, *C. auris* (Antinori et al. 2016; Kullberg and Arendrup 2015; Lamoth et al. 2018; McCarty and Pappas 2016; McCarty et al. 2021). *C. auris* is of particular concern - first described in 2009 following its isolation from a hospitalized patient in Japan (Satoh et al. 2009), it spread globally in less than a decade (Calvo et al. 2016; Chowdhary et al. 2013; Chowdhary et al. 2017; Lee et al. 2011; Magobo et al. 2014). Drug resistance is especially problematic for *C. auris*, resulting in the U.S. Centers for Disease Control recently classifying this fungus as an urgent antimicrobial-resistance threat due to its common resistance to multiple antifungal drugs, ability to spread easily via skin infection in nosocomial settings, and the severity of infections (CDC 2023; see Eix and Nett 2025 for review).

With an urgent need to better understand the biology, pathology, and drug resistance of *Candida* spp., it is essential that our understanding of the experimental data for these species be distilled and synthesized in a single location, so that researchers can access what they need to best design and interpret their experiments. A permanent site such as CGD obviates the need for individual research groups to maintain access to their own tools or datasets. The Candida Genome Database will adapt to meet evolving challenges so that researchers have access to the best information available.

## A Modern Interface and an API for CGD

The CGD project was started April 1, 2004 (Arnaud et al. 2005) and adopted and modified the available codebase (so that it could store data for multiple species) from the SGD project (PMID 39530598), bringing the database online soon thereafter. SGD’s codebase itself had been rewritten over the preceding few years, almost exclusively in the Perl programming language, using CGI and an Apache webserver on the front end to deliver content to the webpage, and an Oracle database on the back end to store the data. The resulting HTML was relatively simple, and modern web technologies, which enable webpages to be dynamic, were not in wide use.

While that code and the pages that were produced have served CGD and its community well for over 20 years, they also limited how we could serve data to our users. We had thus proposed in our most recent grant renewal to replace and modernize the entire software stack with a new codebase, either based on SGD’s current codebase, which has continued to be developed, or from scratch. This would allow us to take advantage of modern web technologies to provide users a more dynamic experience, and to better architect the codebase, with a clear goal of separating the front and back ends.

When we proposed to re-architect CGD, we expected it would be a multi-step, possibly multiyear project, during which we would piecemeal replace CGD’s code. However, between proposing modernizing CGD’s codebase and being ready to do so, the emergence of generative AI greatly simplified and accelerated the project. Using Claude Code, we were able to completely replace CGD’s code in about 4 months. CGD now has cleanly separated back and front ends. The back end uses FastAPI (https://fastapi.tiangolo.com/) and SQLAlchemy (https://www.sqlalchemy.org/) to retrieve data from the database, while the front end uses React (https://react.dev/) and Vite (https://vite.dev/) to present those data to the user. The backend allows us to provide endpoints, which produce JSON responses, that in turn are consumed by the front end to render the display using modern web technologies. This clean separation allows users to also access the endpoints to retrieve JSON data if they want to programmatically access CGD data via our developer API (https://www.candidagenome.org/developer/api).

## Expansion of CGD species to include curation of *C. tropicalis*

Prior to the submission of our last grant renewal in 2024, we surveyed our users to ask which *Candida* species we should incorporate next into CGD for curation. The strongest support was for *C. tropicalis*, which represents 3–66% of all Candida bloodstream isolates worldwide (Pfaller et al. 2000; Sabino et al. 2010; Sipsas et al. 2009; Tan et al. 2010; Viscoli et al. 1999; Yang et al. 2010; Yap et al. 2009) and in many centers is the second most common *Candida* species isolated (see Chai et al. 2010). We have incorporated a chromosome-level gapless genome assembly of the *C. tropicalis* type strain MYA-3404 that was generated from PacBio reads combined with 3C-Seq data (Guin et al. 2020), with gene annotations contributed by Richard Bennett’s lab. Each gene in *C. tropicalis* now has its own locus page, and as we curate the literature, genes with experimental annotations will gain content. We will first curate the literature backlog and then curate new publications in real time as they are added to PubMed.

## New Tools at CGD

### Synteny Viewer

Orthology assignments between related species are useful for evolutionary analyses, but also for understanding more about an ortholog in a more poorly characterized species. We have thus added to CGD the ability to easily navigate between orthologs of different species. The first way that users can navigate to a different species’ gene page for an ortholog is via a new dropdown menu on the top left of a locus page, which allows a user to select the species of interest. Our second method for users to navigate between orthologs is the new Synteny Viewer (Figure 1), which renders a chromosomal region for a focal gene in the species of interest, and then indicates which genes in the other CGD species are orthologous on the syntenic chromosomal regions. The ortholog mapping used at CGD to allow either switching between gene pages for different species, or underlying the synteny browser, is taken from the mappings available at the Candida Gene Order Browser (CGOB; Fitzpatrick et al. 2010; Maguire et al. 2013). CGOB provides orthologous gene assignments that have been extensively manually curated, based on genomic context (local synteny) as well as sequence similarity, providing a “gold-standard” set of orthologs for evolutionary analysis.

**Figure 1.**
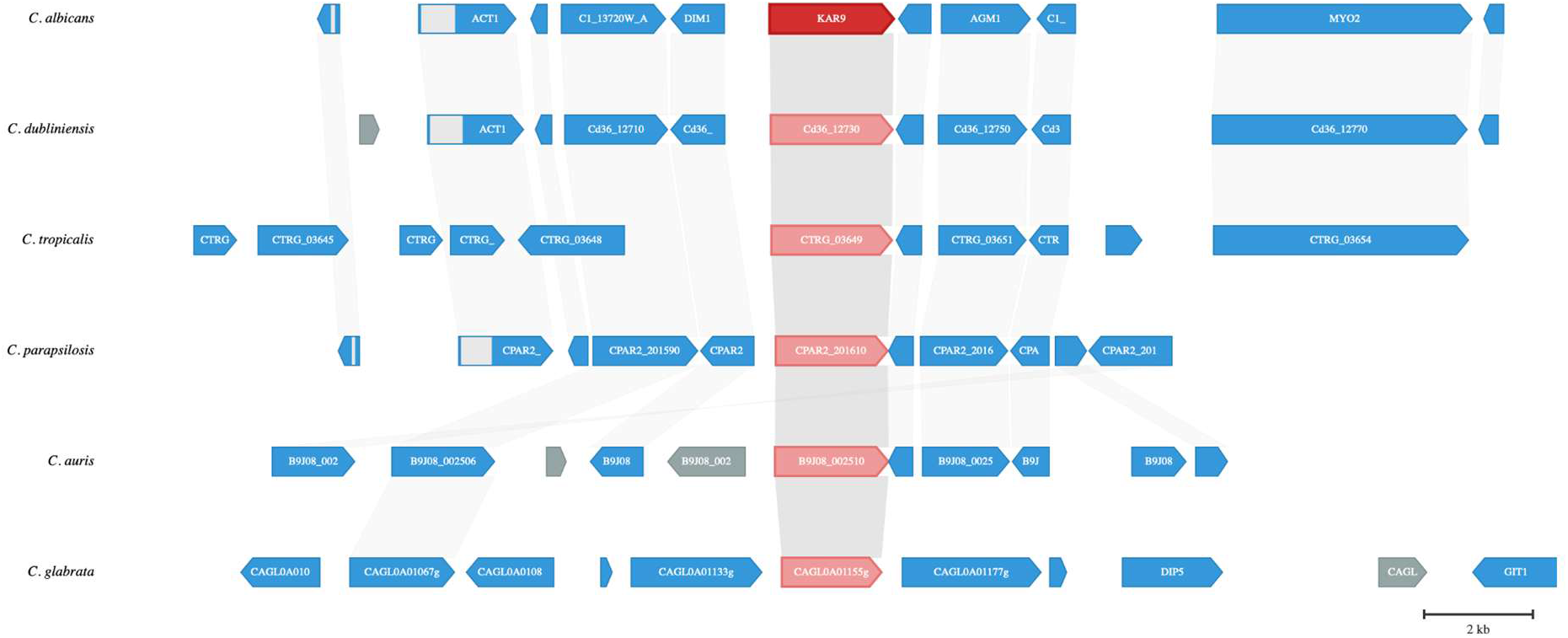
CGD’s Synteny Viewer. The focal gene, *KAR9* in *C. albicans*, is colored red, and its orthologs across the other species are also in red. If a user mouses over genes to the left or right, it will indicate which genes they are orthologous with. Genes colored blue have orthologs but are not in the query. Genes colored gray do not have orthologs among the species set.

### Ortholog Converter

Often when a researcher is working in a *Candida* species, especially one which has not been experimentally well-characterized, it can be challenging to gain biological insights into large-scale data, such as an RNA-Seq dataset. To help users with this challenge, we have implemented a tool that allows the retrieval of a list of orthologs from one species, based on a list of identifiers in the species of interest. For example, a common use is to retrieve the *S. cerevisiae* identifiers for a list of identifiers from a *Candida* species, to take advantage of the extensive gene and protein annotation available from SGD (Engel et al. 2025). As with the Synteny Viewer, we take advantage of ortholog mappings from CGOB (Fitzpatrick et al. 2010; Maguire et al. 2013), as well as the Yeast Gene Order Browser (Byrne and Wolfe 2005).

### Virulence Factor Browser

Because of the importance of *Candida* yeasts as human pathogens, we implemented the Virulence Factor Browser (Figure 2), which allows users to identify *Candida* genes that have evidence linking them to virulence, pathogenesis, or host interaction. It aggregates data from multiple sources, including phenotype annotations, GO terms, and curated literature, to help researchers identify and prioritize genes of interest for virulence studies. We identify genes that have annotations to one or more *Virulence Categories* (Adhesins, Secreted Enzymes, Morphogenesis, Host Interaction, Biofilm Formation, Immune Invasion, and Drug Resistance), and they are further assigned a confidence score with respect to their likelihood of influencing virulence, based on the strength and nature of the supporting evidence (https://www.candidagenome.org/help/virulence-factor-browser). Users can select from the virulence factors in which they have most interest, as well as narrow down genes by species and evidence type, to yield a list of genes that match their criteria. We consider this to be an experimental feature at CGD and encourage users to let us know whether it is useful, or how we might improve the tool.

**Figure 2.**
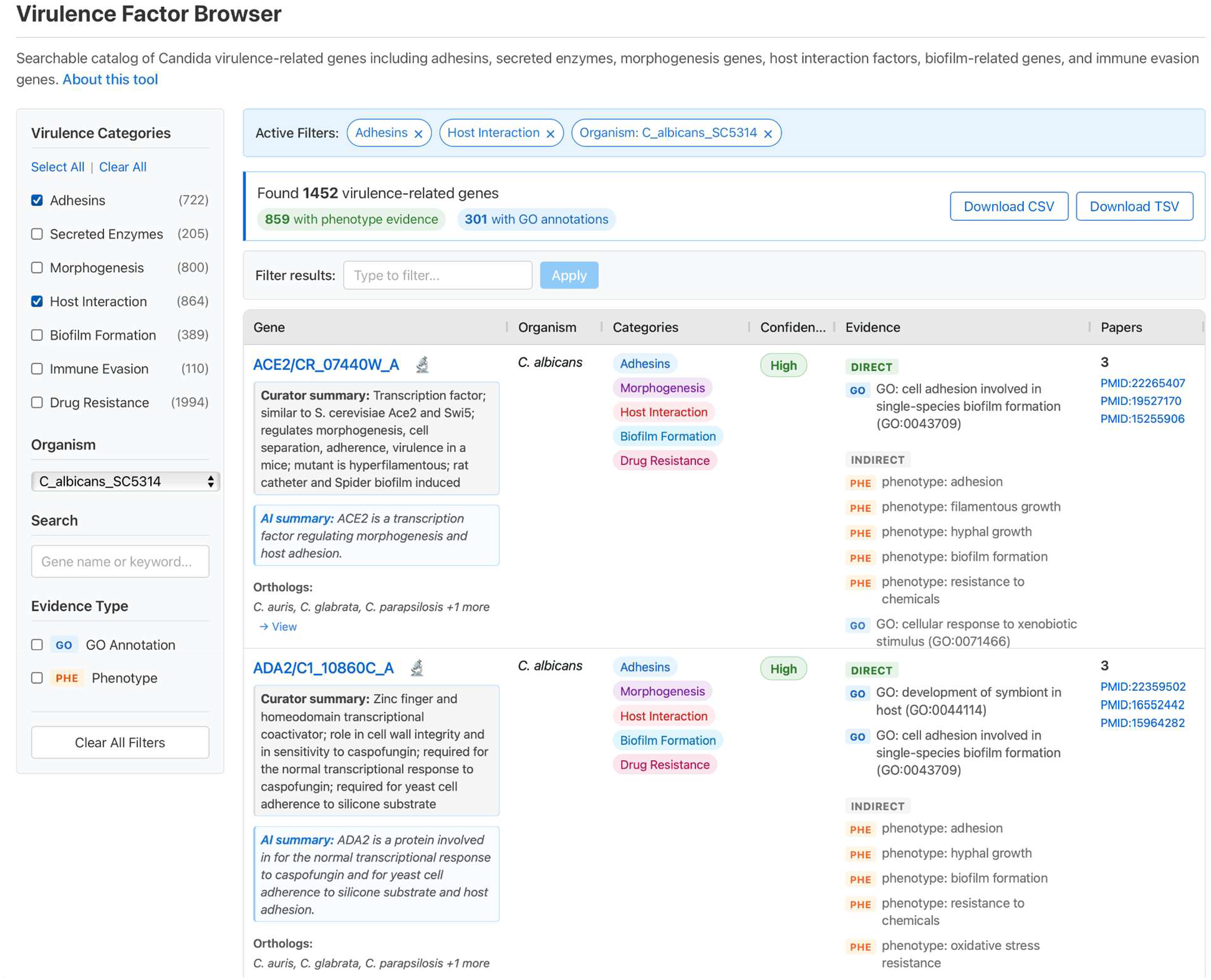
Virulence Factor Browser. The interface includes a curated summary as well as an AI-generated summary, which comprises what is known of virulence-related evidence and/or prediction. The interface also allows for extensive filtering on multiple aspects of virulence data.

### Interaction tab

We have added a new tab to the locus page that displays genetic and physical interaction data for a given feature (Figure 3). These data are sourced from the STRING database (Szklarczyk et al. 2023), BIOGRID (Oughtred et al. 2021) and from annotations curated by CGD. The annotations are provided in list form (with links to evidence), as well as in a Venn diagram, to indicate which sources have identical interaction data, and as a network diagram. Users can pass a list of genes in an interaction network into tools such as GO Term Enrichment. The data from CGD and BIOGRID are manual curations derived from direct experiment, while the STRING database provides powerful computational prediction using data from 12,535 organisms and over 20 billion protein-protein interactions.

**Figure 3.**
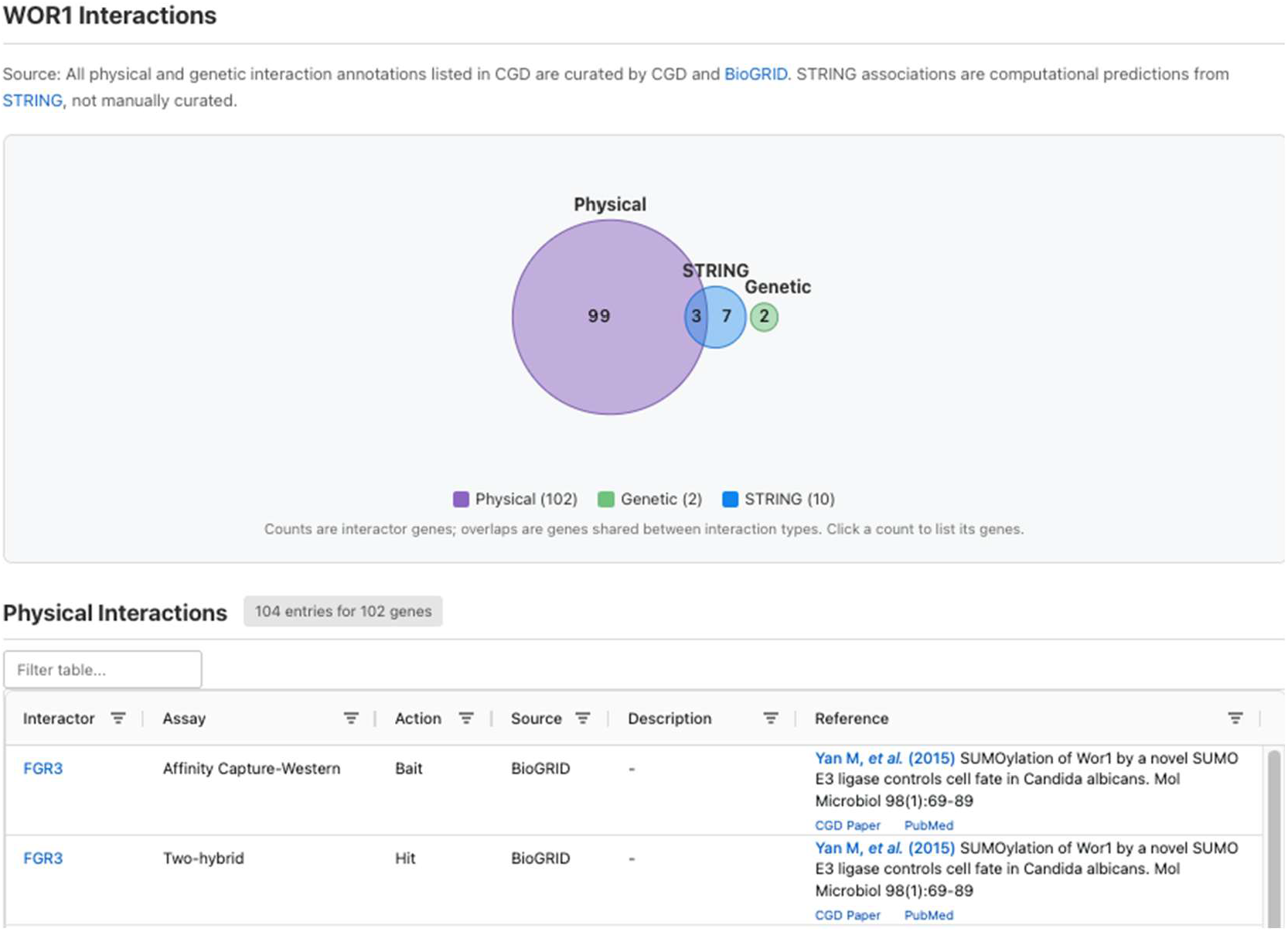
Interaction tab. Interactions search results showing a Venn diagram of physical, genetic, and computationally predicted interaction, along with a table listing each interactor, the type of experiment, and the reference.

The value of these interaction data for researchers is in designing the next set of experimental questions. When a set of experiments determines a gene of interest in one species, the ability to rapidly identify the ortholog in better studied species—together with any interactions defined for that ortholog—provide clues as to which genes to study next in the species of interest. For example, if a mutant screen identifies the previously uncharacterized gene *AAA1* as required for virulence in a poorly studied *Candida* species, use of either the Synteny Viewer or Ortholog Converter tool will identify any orthologs in *C. albicans* and/or *S. cerevisiae*. If the ortholog of *AAA1* in *C. albicans* has been shown to physically interact with *C. albicans BBB1*, while the ortholog of *AAA1* in *S. cerevisiae* is synthetically lethal with *S. cerevisiae CCC1*, the next experiment in the poorly characterized species might be to knock out orthologous *BBB1* and *CCC1* to evaluate the phenotypes related to virulence. Advances in CRISPR-based approaches in *Candida* species (see CRISPR primer design tool below) make knockouts relatively fast even in understudied species.

### Expression tab

At CGD, we continue to collect gene expression datasets as they are published. The challenge is to present these data to users in a format that allows them to understand how a gene is regulated across different conditions, and which other genes are similarly regulated. While we have long since had the ability to view transcript abundance in JBrowse, which allows one to determine if a gene is expressed under a given condition, it does not provide much biological insight. We have thus added an Expression tab to the Locus Page that shows gene expression as a heat map across multiple conditions, allowing users to view genes that have the most correlated expression to the query gene (Figure 4).

**Figure 4.**
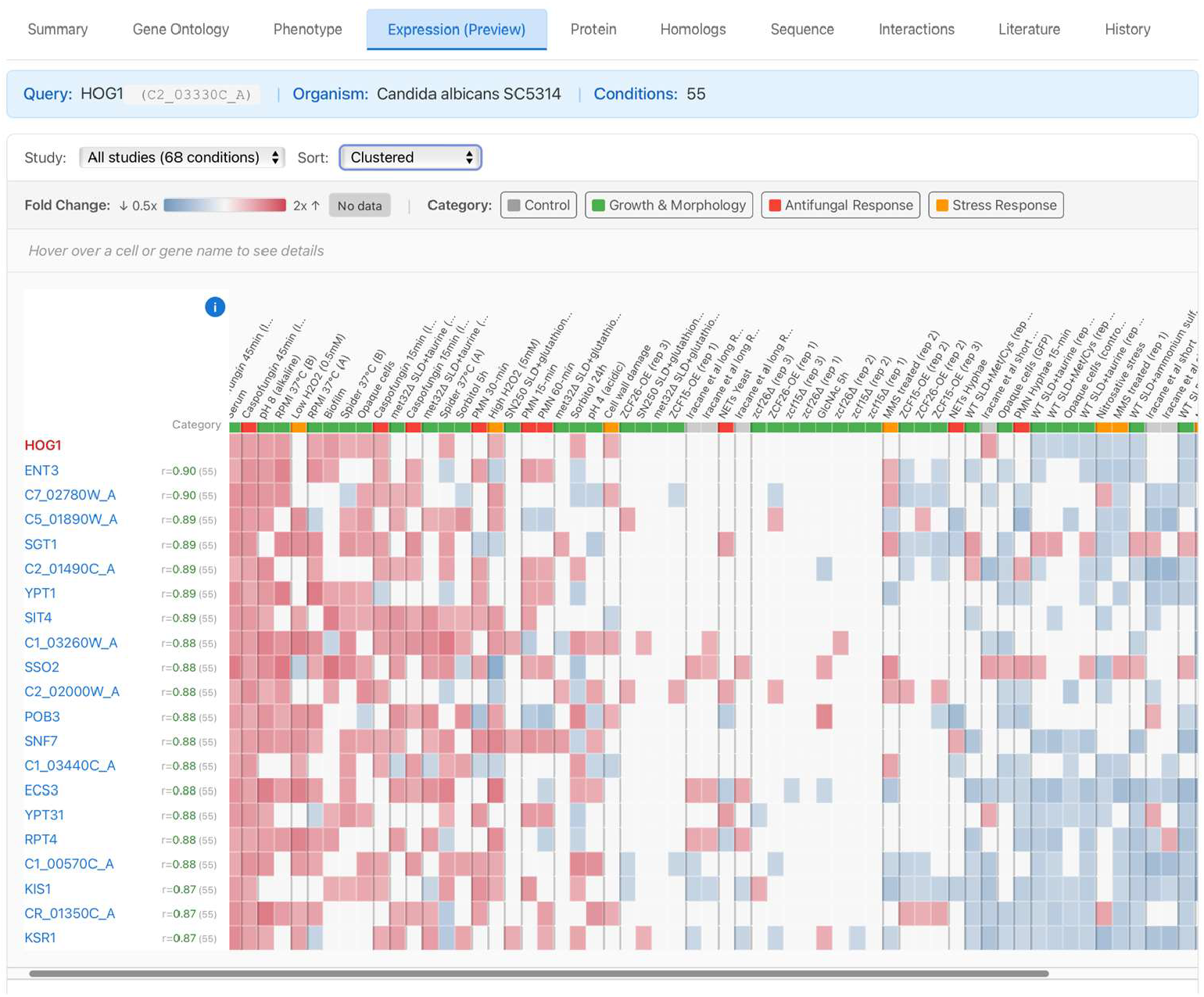
The Expression tab shows a heatmap of gene expression across multiple conditions. Users can sort by the type of experiment conducted to view datasets likely to be relevant to their own research questions. Further, upregulation versus downregulation is highly sortable and searchable.

The genes whose expression is shown in the view can also be passed into other tools, such as GO TermFinder, or to test for phenotype enrichment.

### CRISPR Guide RNA Design tool

CGD recently released the CRISPR Guide RNA Design tool for designing single guide RNAs (sgRNAs) for CRISPR-Cas gene editing experiments in *Candida* species (Figure 5). The tool identifies potential guide sequences within a target gene or DNA sequence, predicts their on-target efficiency, searches for potential off-target sites across the genome, and generates cloning primers for common CRISPR vectors.

**Figure 5.**
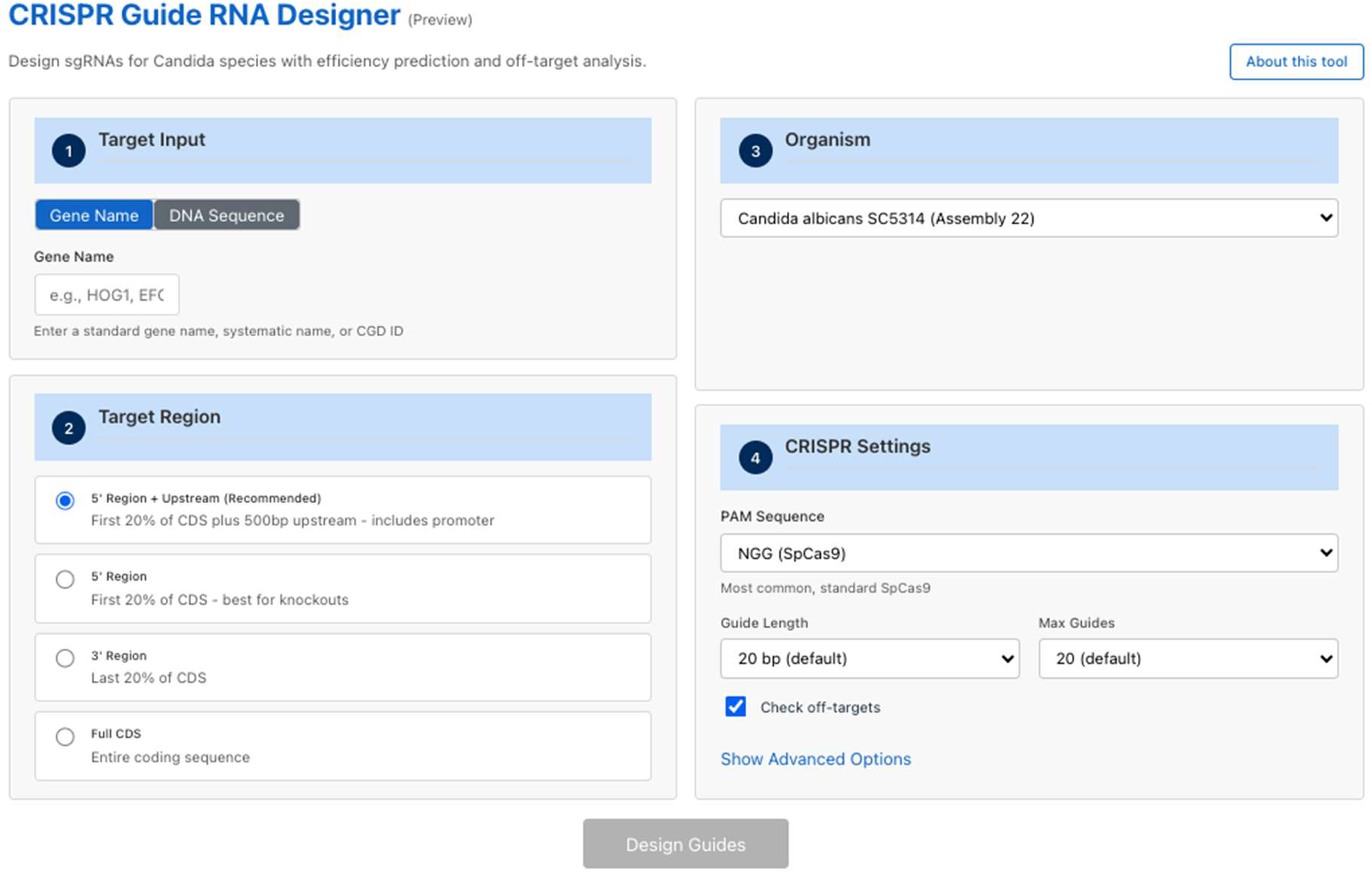
The CRISPR Guide RNA Design Tool makes it straightforward to design high-quality primers for Candida research

The tool is designed to help researchers quickly identify high-quality guides for gene knockout, knockdown, or modification experiments. It is a conservative design tool, in that the default settings allow for targeting only the first 20% of the coding sequence (many tools target the first 50%). While this setting can be changed, the conservative value does the following:

- **Maximizes knockout effectiveness**, where frameshifts near the start codon eliminate more protein function
- **Avoids internal start codons**, where cuts in the first 20% are less likely to produce truncated but partially functional proteins
- **Reduces off-target risk**, where a smaller search region means fewer candidate guides to evaluate, allowing more thorough off-target analysis

The further advantages of using the design tool within CGD for *Candida* research, rather than other widely-used CRISPR design tools, are multifold:

- CGD maintains the most up-to-date genomic assemblies, meaning users can trust the background sequence over widely used tools with hundreds of genomes that may or may not get updated regularly
- CGD’s recommended default option is the “5’ Region + Upstream”, which includes 20% of the coding region plus 500 bases of upstream region. This option is not available on most other sites.
- As with all CGD tools, our methodology is explained clearly and references are provided so users can evaluate the information for themselves.

## Discussion

CGD has made substantial use of Generative AI to replace our prior codebase and to develop new tools for *Candida* researchers. It is important to note that the engineer working on CGD has 30+ years of software development experience in multiple programming languages, as well as a Ph.D. in biochemistry, so it is not that we are blindly using AI, but rather that we are employing it as a tool to speed up our development cycle. We also develop extensive tests for each new tool, and when bugs come to light (which can and does happen) we develop extensive additional tests after fixing the bug, to ensure that the bug cannot recur. Historically, writing code has been the bottleneck to either updating or creating new tools, but with the use of Generative AI, that is no longer the case. Instead, developing ideas and specifications for tools for the *Candida* research community and determining which will be the most useful is the most time-consuming component. Once we have such ideas, implementation is then fairly straightforward, and we invite the community to provide us with suggestions for additional tools.

The future addition of more pathogenic *Candida* species to CGD will depend in part on the scale of the available curatable literature, as the true value of having a species in CGD lies in having aggregated experimental data and annotations in one location. Adding a largely unstudied species to CGD results in numerous blank pages, though the ability to easily switch to an ortholog’s locus page makes this more useful than it otherwise would have been. CGD remains open to the needs of the research community and will consider all requests for additional species.

A likely future addition is AI-generated locus summaries, especially for species for which there is currently little experimental data available; such summaries can be generated by considering what is known about the function of orthologs and other homologs in well-annotated species.

CGD’s manually curated summaries will continue to highlight experimental results from the literature, whereas AI-generated summaries will be able to make predictions for functions and roles that might not yet have been experimentally tested, thus providing context for what is known. We will always make clear which summaries are written by humans and which by AI, so that users can make their own evaluation of the content. Such side-by-side summaries will complement one another to provide a more comprehensive overview for each locus.

Another topic of discussion among databases is the concept of AI-assisted curation. For example, CGD recently used AI to triage the older literature for a species new to the database. AI workflows are excellent at scanning for genes associated with species names among reams of text. By running these searches and then narrowing the hits in various ways, the number of papers required to receive manual curation was straightforwardly narrowed down to a manageable number.

In sum, CGD remains committed to serving as a resource for an important research community targeting a set of dangerous human pathogens.

## Funding

Funding for CGD is providing by R01 DE015873 from the NIDCR/NIH to G.S.

## Conflict of Interest

The authors declare no conflicts of interest

